# Simultaneous Cardiac and Neurological Monitoring to Assess Chemical Exposures and Drug Toxicity in Xenopus Laevis

**DOI:** 10.1101/2021.09.07.459337

**Authors:** Xing Xia, Jimmy Zhang, Manoj Vishwanath, Sadaf Sarafan, Ramses Seferino Trigo Torres, Tai Le, Michael P.H. Lau, Anh H. Nguyen, Hung Cao

## Abstract

Simultaneous monitoring of electrocardiogram (ECG) and electroencephalogram (EEG) under chemical exposure requires innovative engineering techniques that can capture minute physiological changes in studied animal models. However, this is often administered with a bulky system that may cause signal distortions and discomfort for animals. We develop an integrated bioelectronic sensing system to provide simultaneous ECG and EEG assessment in real-time under chemical exposure for *Xenopus laevis*. The microelectrode array (MEA) membrane with integrated ECG and EEG sensing offers an opportunity to achieve multichannel noninvasive electrophysiological monitoring with favorable dimensions and spatial resolution. To validate the performance of our system, we assessed the ECG and EEG of Xenopus under exposure of Pentylenetetrazol (PTZ), an epilepsy-inducing drug. Effects of PTZ were detected with clear ECG and EEG alterations, including frequent ictal and interictal EEG events, 30 dB average EEG amplitude elevations, abnormal ECG morphology, and heart rate changes. Overall, our Xenopus-based real-time electrophysiology monitoring system holds high potential for many applications in drug screening and remote environmental toxicity monitoring.

## 1. Introduction

Pharmaceutical development involves a complicated process of biochemical assays and validation processes in animal models and humans before distribution to market (Barros et al. 2008; MacRae and Peterson 2015; Zon and Peterson 2005). Conventionally, mammalian models for preclinical toxicity, safety assessment, and side-effect screening needed to be determined first before experiments could be conducted. Large mammalian models, such as monkeys and dogs, were used as they possess comparable size and excellent gene similarity to humans. However, they were not always the ideal choice due to high cost, time consumption, and complications in conducting the screens (Denayer et al. 2014). Some mammalian animals, especially dogs and nonhuman primates, faced strong ethical issues for such preclinical tests (Baumans 2004). Hence, small mammalian animal models, such as rodents and rabbits, had been widely used. Although they shared a number of electrophysiological similarities to humans, there were some notable disparities. For instance, their heart rate (HR) was several hundred beats per minute (bpm) while the respective value for humans was ∼60-70 bpm, questioning their suitability for studying the changes in cardiac electrophysiology under the influence of drugs. Recently, other models, such as zebrafish, had been explored, owing to their fecundity, morphological and physiological similarity to mammals, and the complexity of the circadian clock in relation to behavioral, sleep cycle, cellular and molecular responses. For example, in (Cao et al. 2014) and (Lee et al. 2020), ECG and EEG alterations were evaluated for cryo-injury assessment and epilepsy studies. Nonetheless, due to the small size of zebrafish, developing high spatial resolution electrophysiology recording devices for long term monitoring is still challenging. Furthermore, zebrafish do not possess lungs, which limits numerous studies involving the respiratory system, such as COVID-19. All these call for an alternative model that can facilitate drug screening and discovery.

Simultaneous EEG and ECG monitoring had been used to identify neurocardiac dysfunction in human and mouse models of epilepsy (Kendirli et al. 2014; Mishra et al. 2018). As EEG and ECG recording were commonly acquired from different areas on the body, simultaneous monitoring was often implemented separately using two different systems (Seitsonen et al. 2000). As a result, it was challenging to reduce the device size and bring comfort to test subjects. Alternatively, *Xenopus laevis (X. laevis)* has a unique body structure that could facilitate non-invasive recording of electrophysiological signals from the brain and heart. First, *X. laevis’* hairless and highly conductive skin alleviates the difficulties in obtaining high signal to noise ratio (SNR) and stable biopotentials. Second, anatomical features such as the favorable location of the heart, the absence of ribs, and the special structure of the transverse process on vertebrae allows easier acquisition of the posterior ECG than the anterior ECG, which enables the recording of ECG and EEG from the same side of Xenopus body. Third, the short distance between Xenopus heart and brain permits the implementation of electrodes on a single piece of flexible membrane to obtain ECG and EEG signals simultaneously. These advantages provide the opportunity to minimize the device size without sacrificing comfort and signal quality, promoting the use of Xenopus as an alternative model for drug screening studies.

Pentylenetetrazol (PTZ) is an antagonist of gamma-aminobutyric acid (GABA) on GABA receptors via the Tert Butylbicyclophosphorothioate site (Huang et al. 2001). The PTZ compound was used as a typical model for screening potential novel antiepileptic drugs and has become the approved drug (Löscher 2011). PTZ can penetrate through the plasma membrane, allowing PTZ to induce seizuresvia intracellular sites of action (Bloms-Funke et al. 1996). The EEG alterations caused by PTZ effects had been studied in zebrafish (Cho et al. 2017) and *X. laevis* tadpoles (Hewapathirane et al. 2008). Both works reported detections of high amplitude spikes, the criteria for epileptic events. In (Cho et al. 2017), 4 channel EEG was obtained from zebrafish by an EEG electrode array made with gold deposited on a flexible polyimide substrate. However, due to the limited size of zebrafish brain, the electrode size must be deliberated in order to guarantee high quality and localized EEG signals. In addition, careful immobilization and perfusion were needed during recordings as zebrafish and tadpoles cannot stay out of water for long time, increasing the complexity of obtaining EEG signals.

In this work, we developed a single flexible MEA with gold electrodes on a flexible polyimide film to simultaneously assess ECG and EEG in adult *X. laevis*. The acquired multichannel electrophysiological signals were amplified by a differential amplifier and digitized by an analog to digital converter for further data analysis. Using our system, EEG and ECG signals acquired from PTZ-induced seizures were simultaneous monitored, displaying distinguishable changes during ictal and interictal stages as well as their strong correlation over time. The system rapidly and accurately responded to phenotype changes, which offer potential and practicality to build compact wearable electrophysiology monitoring devices based on *X. laevis* for various biological studies and monitoring.

## 2. Materials and Methods

### 2.1. Animals

All animal protocols were reviewed and approved by the Institutional Animal Care and Use Committee (IACUC) protocol (#AUP-21-066, University of California, Irvine). Xenopus were maintained in separated aquaria with 21°C freshwater cycling. The aquaria were placed in a room with a 12 h:12 h light: dark cycle. 29 adult wild-type female Xenopus were used for this study. Due to the distinct differences in body sizes and weight between female and male *X. laevis* and the limited availability of male *X. laevis* in our facility, only female Xenopus were utilized in the experiments. There should be no significant difference among male and female *X. laevis* subjects in terms of electrophysiological assessment.

### 2.2. Microelectrode array (MEA) design and fabrication

To ensure the biocompatibility and flexibility of the sensors for a uniform and secure contact with the skin of Xenopus, we chose to fabricate the electrodes on polyimide films with various thicknesses ranging from 25 to 125 µm (Kapton, Dupont, Wilmington, DE, USA). First, a Au/Cr layer of 200/20 nm was deposited on the polyimide substrate Conventional lithography and wet etching processes were used to pattern the four 50-μm-diameter circular electrodes. After the cleaning and post baking process, the traces of the electrodes were encapsulated by a layer of hardened photoresist, with the electrode areas and contact pads exposed.

In compliance with the brain structure of a Xenopus, we positioned the 4-channel EEG working electrodes (WE) on the scalp above the right and left sides of the telencephalon and the mesencephalon (Fan et al. 2018) (**Fig 1A**). The reference electrode was located above the center of the cerebellum. Three total lengths (25, 35, and 45 mm) of MEAs were designed to cater to different demands of flexibility. The ECG working electrodes were integrated on the MEAs, which have a total length longer than 35 mm. The distance between the ECG working electrode and the reference electrode was 20 mm, referring to the average heart-brain distance of Xenopus we measured. The ECG and EEG recording electrodes share a common reference electrode. Since EEG signals have lower amplitudes than ECG signals, the baseline noises caused by brain signals from cerebellum area were filtered out by additional signal processing. The electrodes were led to contact pads with a size of 2 mm × 0.35 mm each to fit Flat Flexible Cable (FFC) connectors. A subsidiary printed circuit board (PCB) was designed to facilitate better connections between the electrodes and the cables leading to the subsequent signal collecting system. On the PCB, a FFC connector (5034800, Molex, Lisle, IL) with a 0.5 mm pitch was routed to a 2 row, 2.54 mm female socket header connector. The fabricated flexible electrode membrane was easily slipped and locked into a FFC connector, and the signal transmitting cables were connected to the female sockets on PCB by male pin headers. The PCB with connected flexible MEAs were shown in **Fig. 1B**.

**Fig.1.**
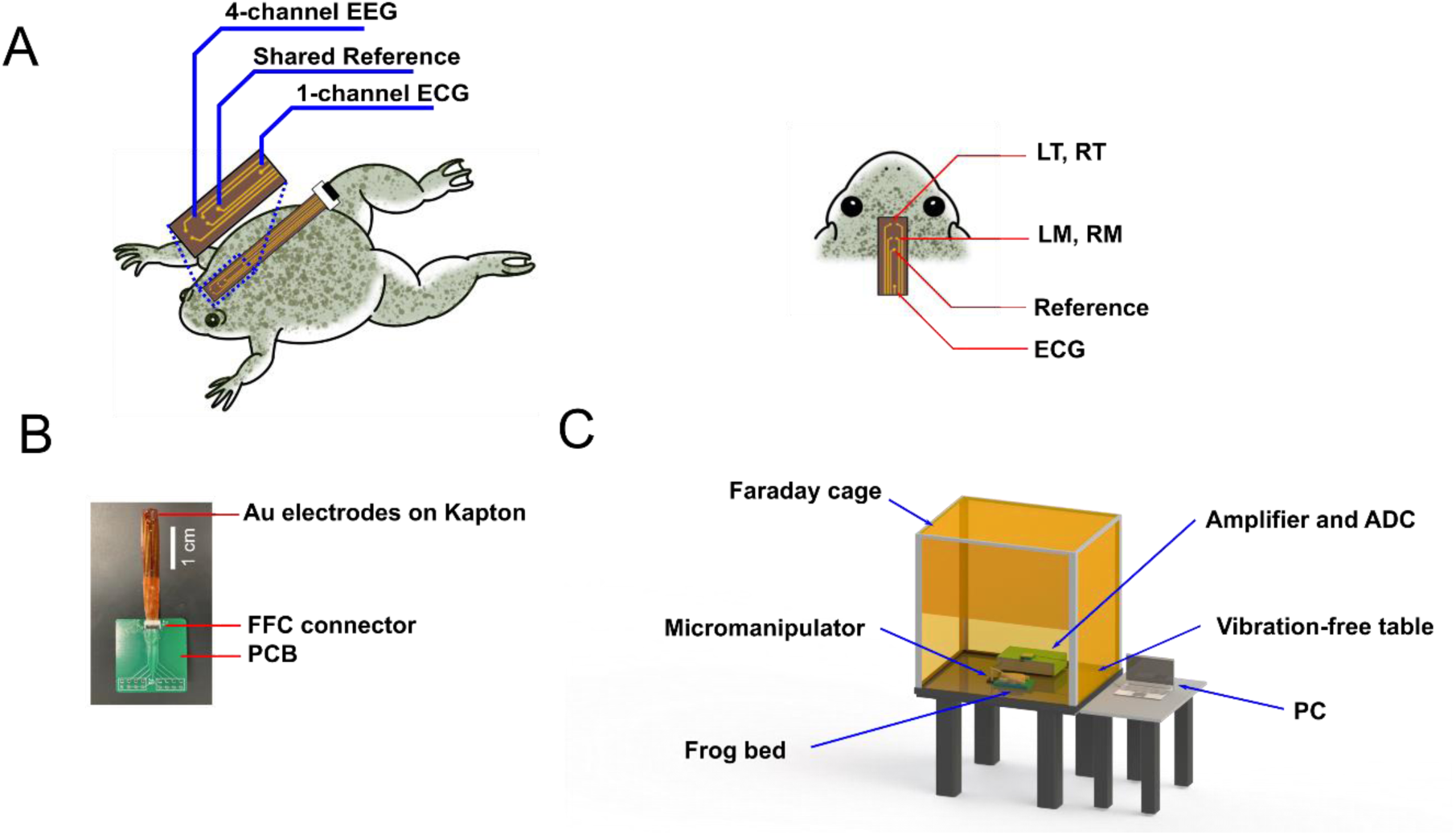
The experimental setup and devices to record EEG and ECG simultaneously. **A**. 4 channel EEG and 1 channel ECG electrodes integrated on one polyimide membrane and the locations of electrodes, including left and right telencephalon (LT and RT) electrodes, left and right mesencephalon (LM and RM) electrodes, a reference electrode, and an ECG electrode. **B**. A picture of the microelectrode array and the PCB connector. **C**. The experimental setup includes a Faraday cage, a differential amplifier, an analog-to-digital converter, and a laptop for data acquisition.

### 2.3. ECG and EEG signal acquisition

*X. laevis* subjects were deeply anesthetized by immersion in a buffered solution of 1.5 g/L tricaine methane sulfonate (MS-222). The subjects were considered sufficiently anesthetized with the loss of toe pinch response. After anesthesia, the Xenopus was settled on a specially designed frog bed. The test bed was made with polydimethylsiloxane (PDMS), and possessed a concave surface to fit the shape and flexibility of frogs’ abdomens. Then, the MEA was placed dorsally based on the location of brain and heart and fixed by a manual micromanipulator (M3301, World precision instruments Inc., Sarasota, FL) to reduce vibrations and noises. Due to the different shrinkages of the frog skin and the Kapton membrane, the electrodes automatically adhered to the skin resulting in a better electrical connectivity. During the entire experiment, the anesthetized Xenopus and accompanying recording devices were placed in a Faraday cage on a vibration-free table. The signals were amplified 10,000-fold by a voltage differential amplifier (A-M Systems Inc. 1700 Differential Amplifier, Carlsberg, WA) and filtered between 1 and 500 Hz with a cut-off frequency of 60 Hz (notch). The filtered signals were then digitized at a sampling rate of 1,000 Hz using a Data Acquisition Hardware (National Instruments USB-6251 DAQ device, Austin TX, and LabVIEW), and stored in a laptop for further data analysis. The full system setup is sketched and shown in **Fig. 1C**.

### 2.4. Drug exposure

To validate the functionality of the system, we conducted PTZ-induced epilepsy and anti-epileptic drug tests using valproic acid (VPA). Due to the excellent absorbability of Xenopus skin, direct immersion into drug solvent was utilized to treat the animal. The Xenopus were placed into a container filled with 200 mL drug solvent, with their eyes and noses above water. The drugs were transported in passively. Compared with injection, this method took a longer duration for the drugs to take effect. 30-mM PTZ dissolved in 1.5 g/L MS-222 was used for PTZ drug exposure. The Xenopus were immersed in PTZ solution for 15 minutes. For the VPA treatment, the Xenopus were immersed in 3-mM VPA solution for 1 hour. During the drug treatment, the containers were covered by lids to prevent the rare case of Xenopus escaping the containers. After recordings, Xenopus were moved to a tank with flushing water to help them recover from anesthesia and drug effects. It took up to 30 minutes for Xenopus to fully recover. The drug-treated Xenopus were separated and maintained in recirculating tanks for 7 days.

### 2.5. Data processing

The ECG signal consisted of certain characteristics such as the P wave, PR interval, QRS complex, ST segment, T wave, and QT interval. For accurate detection of these characteristics, clear ECG signal was required. Therefore, noise cancellation was critical and essential. The lowpass filter and Savitzky-Golay finite impulse response (FIR) smoothing filter (SG filter) were used to suppress the powerline interference noise and remove the baseline wander. The baseline drifted due to respiration and could be normalized by using the lowpass filter. The SG filter proposed by Savitzky and Golay (Awal et al. 2011; Savitzky and Golay 1964) was based on local least-squares polynomial approximation; and it was demonstrated as one of the most popular noise cancellation and smoothing methods.

The obtained EEG signals were filtered using a 6^th^ order Butterworth bandpass filter from 0.5 – 40 Hz. The selection of this frequency range was based on the frequency bands of interest, which includes delta (0.5 -4 Hz), theta (4 – 8 Hz), alpha (8 – 12 Hz), beta (12 – 30 Hz) and gamma (> 30 Hz). The Fourier transform (FFT) of the filtered signal was then calculated and visually compared between different groups to analyze the difference in frequency compositions of the signals.

## 3. Results

### 3.1. Xenopus EEG and ECG recording

Five animals were used for device validation and then 24 *X. laevis* were divided into 4 groups for the drug-induced epilepsy study. Each Xenopus frog was treated with drugs, and performed their 4-channel EEG and ECG recording were obtained 3 separate times. After each experiment, Xenopus were returned to circulating fresh water and allowed to recover from drug or anesthesia effects for at least 2 days. Xenopus in different groups were raised in different aquaria. Each recording lasted 20-30 minutes, depending on the duration that Xenopus were in anesthesia. It took approximately 12 minutes for Xenopus to be fully anesthetized in 1.5 g/L MS-222 solution. Whenever the Xenopus lost toe-pinch reactions, we transferred the anesthetized Xenopus from the anesthesia solution to the frog bed. Adjusting the position of the Xenopus and adjusting the position of the MEA membrane took 2-3 minutes. Three different lengths (25 mm, 35 mm, 45 mm) and 3 different thicknesses (25 μm, 75 μm, 125 μm) of the polyimide membrane were tested. The 75-μm thick, 25-mm long probe and the 125-μm thick, 35-mm and 45-mm long probes displayed the best performances. These probes had similar flexibility and strength to the skin of Xenopus; thus, the conformability and longevity were significantly stronger, especially for long term recordings. In some of the recordings, we found that the EEG signals were noisy if the Xenopus scalp had too much mucus. We used paper towels to gently wipe the area before applying electrodes, greatly improving the signal quality. The use of a vibration-free table and a Faraday cage also contributed to obtain a favorable signal-to-noise ratio (SNR) (Defreitas et al. 2012; Tran et al. 2019).

A typical recording of PTZ-treated Xenopus 4-channel EEG is shown in **Fig 2A**. The signals were obtained from the electrodes placed near the left and right sides of the telencephalon (LT and RT) and the left and right sides of the mesencephalon (LM and RM). This segment of recording included 20 seconds of normal EEG (baseline) followed by 20 seconds of the EEG during seizures of epilepsy. Periodic sharp spikes with much higher amplitude than the baseline EEG were observed and defined as the criteria for ictal seizures. More detailed EEG during seizures is shown in **Fig. 2B**. The signals from the left and right sides of the brain were similar in patterns and amplitudes, but comparing the signals from the telencephalon and the mesencephalon displayed numerous differences. Some spikes were clear in the LM and RM channels but difficult to see in LT and RT channels. Some high spikes were contrary in position, implying the origin of these abnormal electrical signals may be from the midbrain area (Saccomanno et al. 2020; Truszkowski et al. 2016). **Fig. 2C** shows the typical patterns of EEG before and after each seizure. There were periods of signals that had much higher amplitude spikes than regular EEG. We identified these spikes with 200-600 μV high amplitudes as an ictal event. In the 20 seconds of recording shown in **Fig. 2C**, there were two ictal events. These high amplitude spikes appeared 3-5 times in one second, and each ictal event contains 5-10 occurrences of such spikes, consistent through the whole recording. Between the two ictal events, some periodic bumps with lower amplitude and less sharpness were detected less frequently. These bumps were identified as an interictal event. The interictal events only appeared after the first appearance of the ictal event, and they disappeared before the ictal events ended. The small spikes of interictal intervals were around 30 μV, occurring 1-2 times in one second. The ictal and interictal events were not detected in the control group due to absence of focal epilepsy (Zijlmans et al. 2011).

**Fig. 2.**
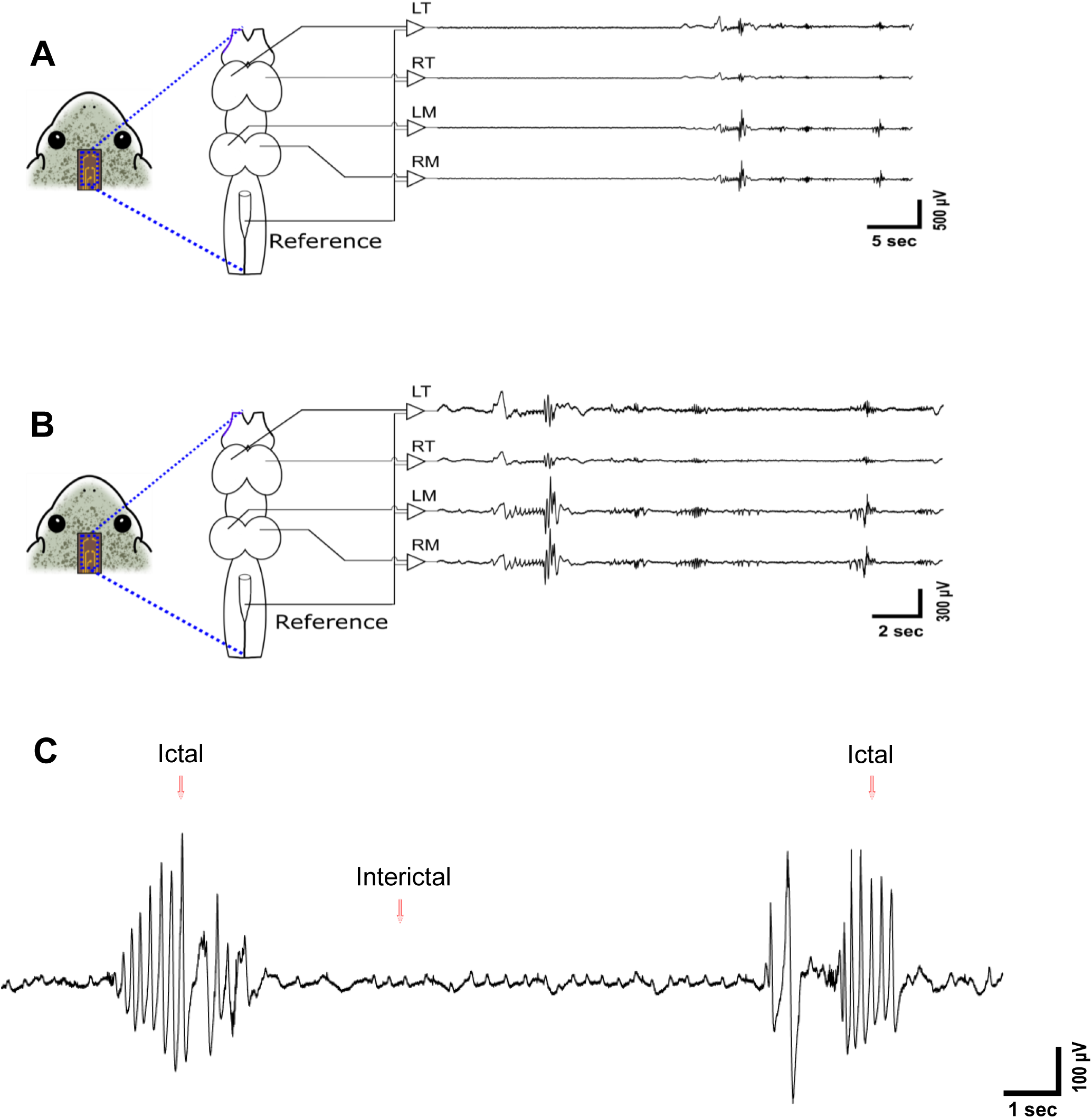
Typical EEG signals recorded by the system. **A**. 4 channel EEG from different locations of the brain in 40 seconds. **B**. 4 channel EEG when a seizure happened in 20 seconds. **C**. Typical EEG patterns of a seizure, includes two ictal events and one interictal period.

The simultaneous ECG signals were recorded by the working electrodes that were dorsally placed close to the heart. We also attempted to obtain ECG signals from the front chest of the Xenopus. Because of the body structure of Xenopus, it did not exhibit as high of a SNR as the ECG obtained posteriorly (Levis 2015). As a result, the ECG recording electrode was placed close to the EEG electrodes. decreasing the size of the MEA membrane to less than 2 cm × 0.5 cm while maintaining high quality ECG and EEG. This result strongly attests to the benefit of using Xenopus as the model for ECG and EEG simultaneous monitoring. To minimize the size of the device and maximize the comfort the animals while monitoring, we chose to use the same reference electrode for both EEG and ECG that was placed on the cerebellum of Xenopus. Due to the short distance between heart and brain of the Xenopus, ECG and EEG signals from cerebellum were obtained with the same set of electrodes (Devuyst et al. 2008; Tong et al. 2001). Despite obtaining mixed ECG and EEG signals from the same channel, the recorded ECG signals were good enough to be used for morphological analysis. **Fig. 3A** shows an example of a portion of the raw signals recorded by the ECG electrodes. The R peaks had amplitudes around 250 μV, which were much higher compared with the baseline noises (less than 20 μV). The R peaks were definable without additional filtering, so the heart rate and R-R intervals could be detected from the raw signals manually. We also tried to record the ECG with an individual ECG reference electrode placed on the right leg of Xenopus. While it provided better ECG signals, the trade-off of the total sizes and the signal qualities made it less than ideal. **Fig. 3B** shows the ECG signals after noise cancellation and smoothing filtering. After data processing, the PQRST waves were detected clearly. The normalized ECG of Xenopus with PQRST marked is showed in **Fig. 3C**. We also obtained the human ECG and zebrafish ECG signals using the same system. Compared to human and zebrafish ECG, Xenopus ECG exhibited one extra wave between the S and T waves. This wave could be from PTZ-induced seizure for ECG change particularly in high-susceptibility frog individuals. ECG changes due to PTZ-induced seizure had not been well studied in Xenopus models; however, advantages in cardiac and brain systems of this model allow correlation of electrographic changes in the setting of drug-induced seizures to cardiac arrhythmia (Desai et al. 2017; Kuttab et al. 2019).

**Fig. 3.**
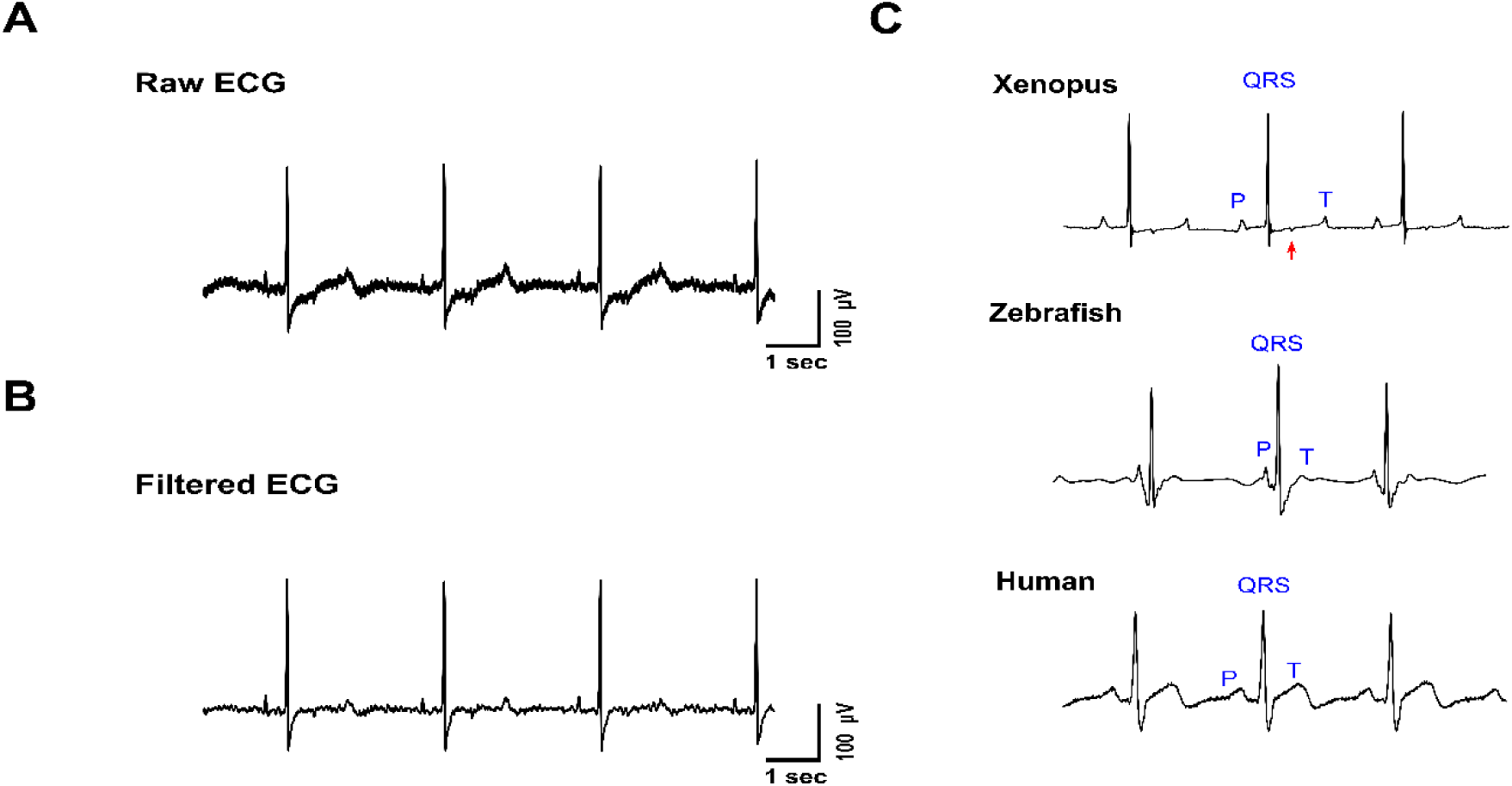
ECG patterns of Xenopus. **A**. The raw ECG signals recorded by the electrodes posteriorly placed. The signals were contaminated by noises and some EEG signals. **B**. The filtered ECG. Now the PQRST complex are more recognizable. **C**. The typical ECG of Xenopus, zebrafish and human recorded by the same system.

### 3.2. PTZ-induced seizures and VPA anti-epileptic effects

The ECG and EEG signals were recorded and analyzed from 4 groups of Xenopus. 12 Xenopus were divided into 4 groups with different drug treatments: 1) PTZ group, in which the Xenopus were treated with 30 mM PTZ mixed with MS-222; 2) VPA+PTZ group, in which they were immersed in 5 mM VPA solution for 1 hour before they were transferred to the same PTZ treatment as the PTZ group; 3) VPA group, in which Xenopus were treated with 5 mM VPA for 1 hour; and 4) Control group, where no drug treatment was administered. Each Xenopus was anesthetized in 1.5 g/L MS-222 and then recorded for more than 25 minutes.

The typical 1500-second-long recordings of raw EEG signals from the control group, PTZ group and the VPA+PTZ group are shown in **Figs. 4A-4C**, respectively. The 1500-second-long samples were recorded by the LT working electrodes (channel 1), but the other 3 EEG channels showed similar results statistically, as all channels were able to record the same number of ictal events. The signals of the experimental groups were significantly different. The most distinguishable feature was the difference in the number of PTZ-induced seizures. EEG recorded in the control group did not show any occurrence of seizures. The average amplitude of control group EEG was 10 μV. For the PTZ group and the VPA+PTZ group, the ictal event occurrence, previously identified by spikes in amplitudes of 200-600 μV, was 34 times and 2 times, respectively. However, upon a closer look into the EEG patterns in the VPA+PTZ group, the epilepsy-like EEG spikes still existed and had lower amplitudes (Smith 2005a). The average amplitudes of the ictal spikes decreased 81.0% from 370 μV to 70 μV. As a result, the criteria of an ictal event for VPA+PTZ group was changed to 37.8-113.5 μV, proportional to the decrease in amplitudes of spikes. After recalculation, the ictal events in the VPA+PTZ group in the 1500-second-long recording appeared 9 times, resulting in a 73.5% reduction in comparison with the PTZ group subject. These results proved that VPA can mitigate PTZ-induced seizures in terms of occurrences and intensities which is in accordance with studies on other models (Puig-Lagunes et al. 2016; Romoli et al. 2019). The recordings obtained from different groups also validate our device for its capability and longevity to collect high SNR EEG for long term monitoring.

**Fig. 4.**
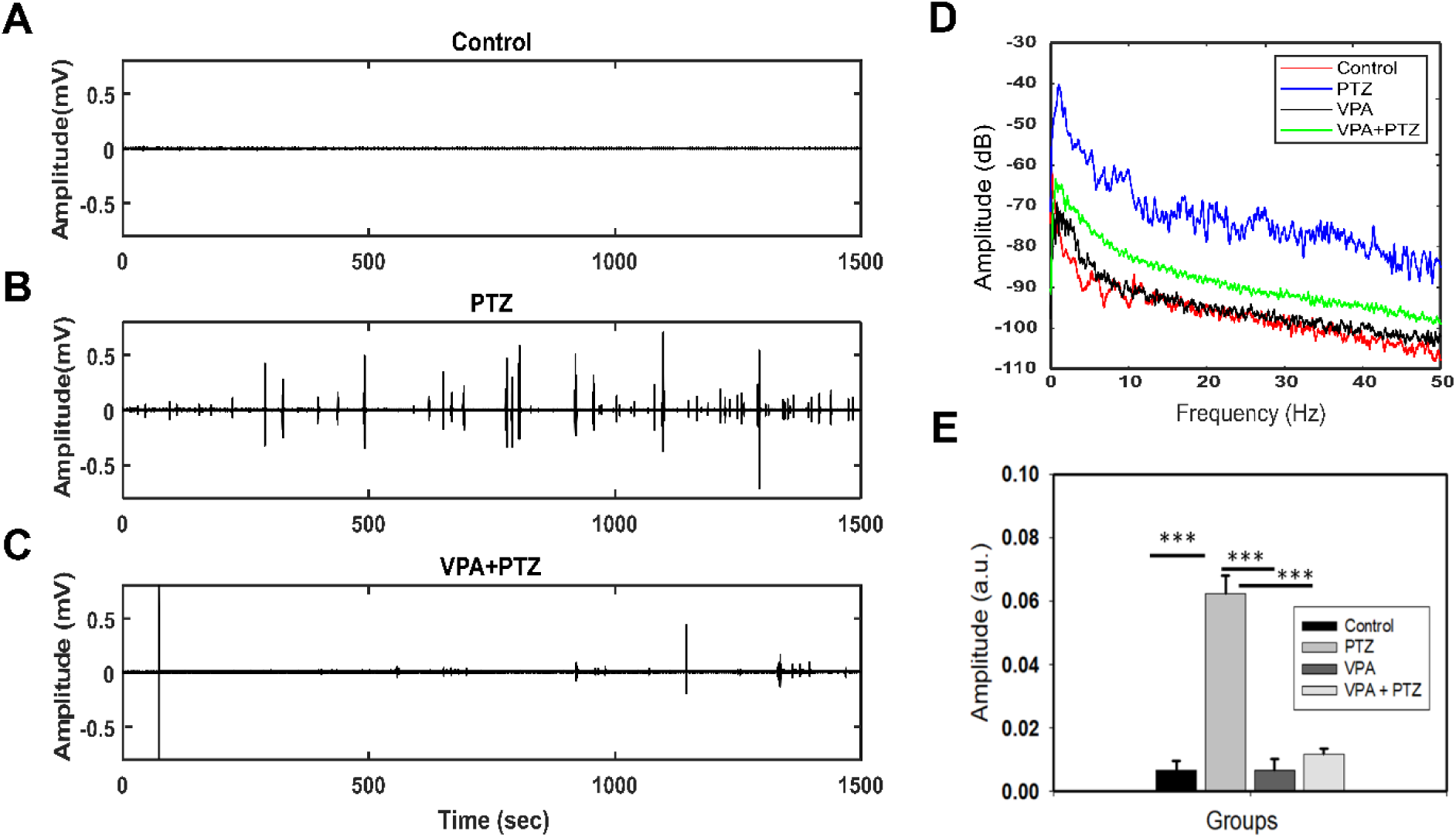
PTZ and VPA effects to Xenopus EEG. **A**. 1500 second EEG of Xenopus in control group; **B**. in PTZ group; **C**. in VPA+PTZ group. **D**. Fast Fourier transform of the EEG in 4 groups. The groups treated with PTZ showed higher amplitude in 0-40 Hz frequency domain, but the VPA inhibited the PTZ effects. **E**. The average amplitude of EEG in 4 groups for quantified comparisons.

The Fast Fourier Transform (FFT) spectrums averaged across 4 recorded channels of the 4 groups are shown in **Fig. 4D**. It quantified the amount of different frequency components present in the given signal. In general, all 4 groups had higher amplitudes in low frequencies, which is a characteristic for all EEG signals. It is observable that the amplitude of the PTZ group was 30 dB higher than the control group and the VPA group for the entire frequency range of frequencies (0-50 Hz). The averaged amplitude of EEG signals in 4 groups over 200-second-long recordings was also calculated and compared. The Control group and the VPA group did not have noticeable differences, indicating that the VPA did not cause significant EEG changes in amplitudes. The results are shown in **Fig. 4E**. With the immersion of PTZ, the average EEG amplitude increased 833.9% in the control group. In the VPA treated groups, the amplitude only increased 70.0% after the immersion of PTZ, resulting in a 91.6% mitigation of PTZ effects. The averaged amplitude in the control group and VPA treated group did not show significant differences, which indicated that the VPA did not have mitigation effects on non-seizure EEG amplitude.

### 3.3. Abnormal ECG and its features caused by PTZ

In our experiments, the heart rates, R-R intervals, QTc intervals and morphological characteristics of Xenopus ECG were analyzed and compared. For the 6 Xenopus frogs in the PTZ group, we calculated the average heart rate every 2 minutes, from the first to the 29^th^ minute mark in the recording. Since the heart rate of Xenopus differed widely in certain individuals (from 11 bpm to 52 bpm in our dataset), the percentage change of heart rate was chosen as the parameter to be analyzed. The average heart rate of the whole 30 minutes recording was calculated individually. The relative heart rate (RHR), which is the heart rate at one moment in time divided by the averaged heart rate of the whole recording, was used to determine the relative variation of heart rate during the whole recording. The average HR variation of the 6 Xenopus after the treatment of PTZ were shown in **Fig. 5A**. To investigate the effect of seizures on the heart rate, we first determined the occurrences of epileptic seizures from simultaneously recorded EEG signals. The first appearance of a typical ictal event in the EEG signals was defined as the boundary of the preictal stage and the ictal stage (Fisher et al. 2014). The boundaries appeared at around 10 minutes into the 6 ECG recordings. The 2 stages were marked with different colors in **Fig. 5A**. By investigating the treads of the Xenopus heart rate variation in PTZ group, we found that the relative heart rate did not increase when seizures occurred. On the contrary, the heart rate was even lower in the Ictal stage than the Preictal stage for all the Xenopus tested. To study the anesthetic inhibiting effects on the heart rate, the control group was were introduced and compared. The control group consisted of Xenopus that were only treated with MS-222 at the same concentration and duration as other experimental groups. **Fig. 5B** showed the relative heart rate (RHR) variation in 30-minute-long recordings of the control group. The RHR curve of the PTZ group was shown in **Fig. 5B** as the red dotted line for comparison. Both groups had very similar heart rate variation from the 10th – 30th minutes of the recording, which was defined as Ictal stage in the PTZ group. But the RHR variation from the beginning to 10th minute in the control group recording was more moderate. This comparison indicated that the PTZ has caused increases in heart rate, but the heart rate gradually returned to normal. Additionally, the ictal events did not have direct relations to the heart rate.

**Fig. 5.**
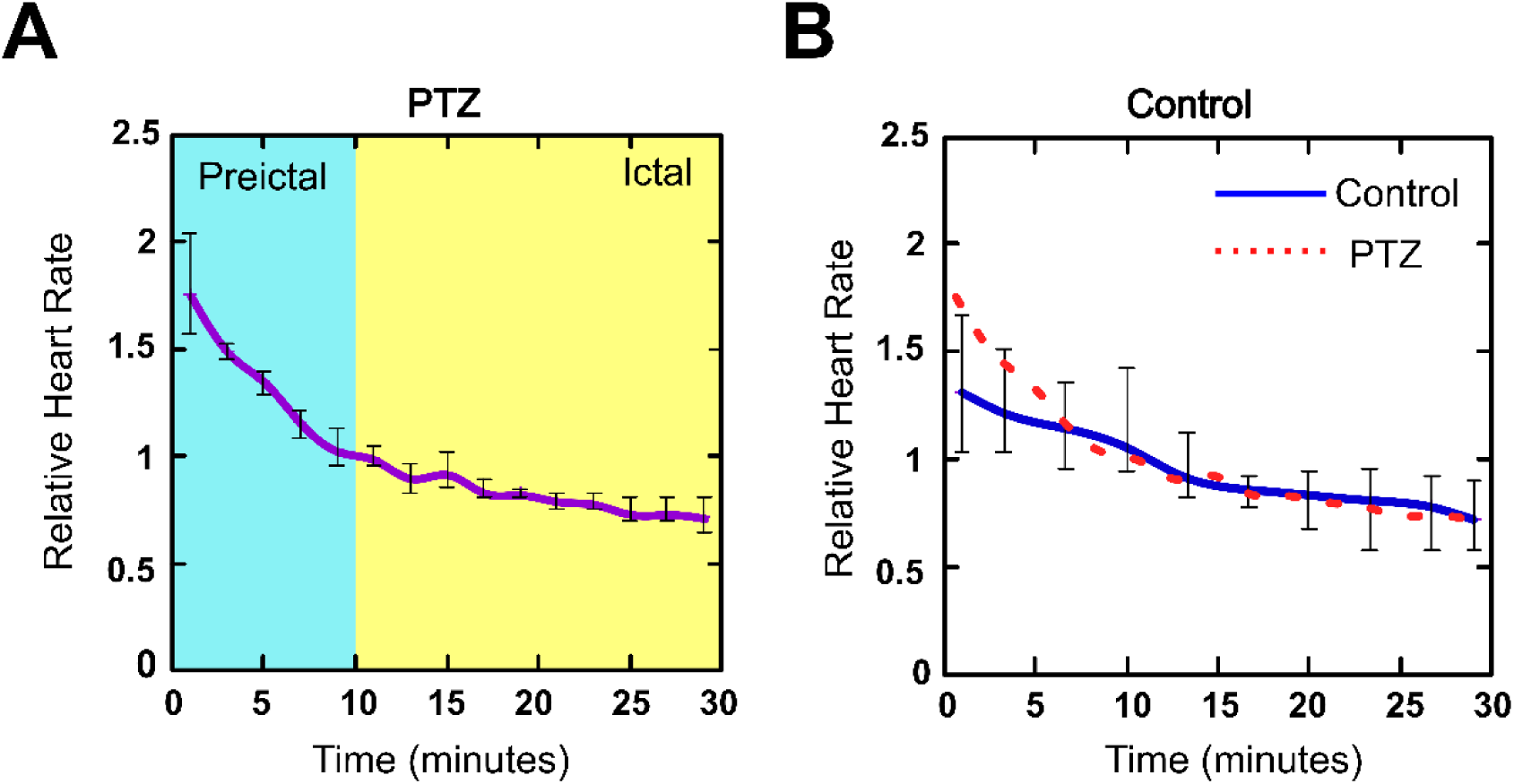
Relative heart rate altering. **A**. The relative heart rate changings during the 30 minutes recordings in PTZ group. **B**. The comparison of relative heart rate trends in PTZ and Control group.

PTZ also induced morphological changes on the ECG patterns (Akyuz et al. 2021; Ruby et al. 2008). The ECG morphology changes were not identical for all Xenopus frogs. From the PTZ group, one Xenopus was found to possess gradually inverted T waves during the recording. The ECG signals after PTZ treatment was shown in **Fig. 6A**. Four intervals of ECG were selected, which are (a), the start of the recording; (b), the end of the preictal period; (c), the start of the ictal events; and (d), the end of the recording. From (a) to (d), the R-R intervals increased from 1.7 seconds to 4.7 seconds, and the T waves changed from upright in (a) and (b), to flat in (c), and then inverted in (d). As the figure depicts, the morphology of other waves did not have significant changes besides T waves. Another Xenopus was detected to have 3 minutes of arrhythmia when the seizures happened. The ECG is shown in **Fig. 6B**. The arrhythmia happened right after the occurrence of the first ictal event. Before and after this 3-minute interval of arrhythmia, the R-R interval was around 2.8 seconds. When the arrythmia occurred, the heart beats were in pairs as one fast heart rhythm and one slow rhythm. The R-R intervals were 1.6 seconds and 2.7 seconds, respectively. In contrast, the morphology of ECG did not have noticeable changes for all subjects in the control group.

**Fig. 6.**
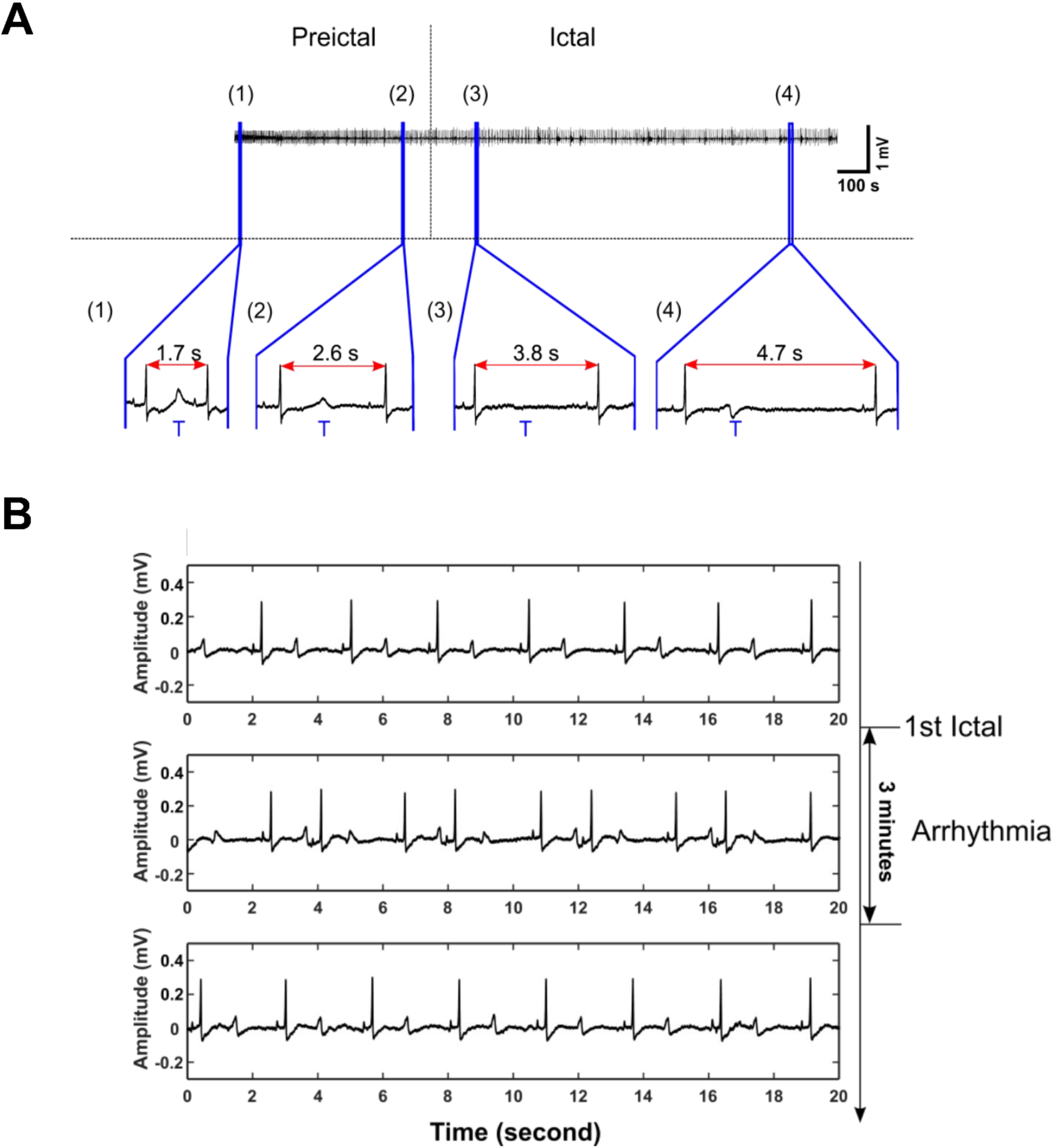
Abnormal ECG found in PTZ-treated Xenopus. **A**. T wave gradual inversion in a 30-minute recording. **B**. 3-minute arrhythmia happened right when ictal events started.

## 4. Discussion

### 4.1. Simultaneous ECG and EEG monitoring

Electrophysiological monitoring is important to biological and medical research. ECG and EEG recordings in humans have been widely used in clinical diagnostics and vital sign monitoring for decades. In recent years, as the development of fabrication technology and electrical engineering, small feature size sensors and electronics were applied into wearable and implantable devices. These efforts provided opportunities to implement high spatial resolution ECG and EEG monitoring on small animal models. Rats and mice were some of the most popular lab animal models. Many researchers presented wired or wireless multichannel rodent EEG recording devices (Choi et al. 2010; Pinnell et al. 2016). Some works also included a simultaneous ECG (Mishra et al. 2018). ECG and EEG recordings on zebrafish were realized as well, but integrating them is still challenging. Many of them used flexible electrodes to reduce the size and discomfort. Nevertheless, most of electrophysiology recording systems were based on rodents or zebrafish. The other animal models were lack of explorations.

During the first part of our work, we tried to place the ECG electrodes on different locations to acquire the best signal. Initially, we considered the chest to be the best location, similar to most other animals. However, we did not obtain ECG signals with satisfying quality, probably because of their ventral fat. Surprisingly, the unique location of Xenopus heart enabled the recording of good ECG signals dorsally. This finding compelled us to combine the ECG and EEG recordings in one piece of sensor for simultaneous recording. Another advantage that the Xenopus offered was their lack of neck structure. This characteristic not only flattened their dorsum but also significantly decreased the distance between the heart and the brain. The special body structure of Xenopus enabled simultaneous ECG and EEG recordings by a set of electrodes fabricated on a small piece of flexible substrate, which offered the opportunity to further minimize the device size.

In our work, non-invasive electrophysiology was chosen. Due to the thin skin and fat layer between the measuring spots and the signal sources, we were able to obtain clear ECG and EEG signals. The drawback of the closed heart and brain was the intermixing of ECG and EEG signals, but it was resolved by signal processing. In the future, invasive electrophysiology with implanted electrodes may be an option as it may provide signals with higher quality and stability.

### 4.2. Cardiac and cerebral influences caused by chemical induced epilepsy

EEG is the most common diagnostic indicator for epilepsy diagnosis. During seizures, abnormal brain waves occur in EEG signals, often manifested as slow wave discharge around 3 Hz (Smith 2005b). High-density EEG was required to precisely determine the affected brain sections and the origins of seizures. However, in small animal models, normal electrodes did not provide enough spatial resolution. Thus, microelectrode arrays needed to be designed and fabricated. In our results, clear slow wave spikes were detected in all channels. It was noticeable that the signals from left mesencephalon were usually the highest in amplitude during seizures. This result indicated the origin of the discharges.

Even though researchers had determined that seizures originated from certain parts of the brain, the cardiac effects of seizures were also widely studied (Bloms-Funke et al. 1996; Huang et al. 2001; Nei 2009). In many clinical observations, the ictal events were linked to an increase of heart rates and many morphological changes (Eggleston et al. 2014). Chemical toxins were possible factors that have been reported to causes epilepsy, seizure or acute clinical syndromes (Jett 2012). The chemicals were toxic to the nervous system by causing hyperactive neurotransmission generated by a temporary change in the electrical functioning of the brain. In seizures, the brain’s electrical rhythms were disrupted by sudden and synchronized bursts of electrical energy which tend to become imbalanced, resulting in alterations of the heart rate and abnormal rhythm (Nei 2009). Thus, we theorized that there was a possible connection between cardiac associated sudden unexpected death and epilepsy. Although the seizure-associated cardiac arrhythmias, such as ictal asystole that causes syncope, have been discovered, the overall risk of seizures on cardiac status still remained elusive. Recently, ECG and blood pressure results associated with PTZ-induced seizures and epilepsy were frequently encountered in elevated catecholamine release, which induce abnormal ECG (Akyuz et al. 2021; Ghosh et al. 2009; Nass et al. 2019; Zavala-Tecuapetla et al. 2020). Unlike most reported human heart rate changes, our results showed that the PTZ induced seizures did not cause an increase of heart rate while ictal events happened. This observation was consistent with a previous report (Randall and Rohse 1956), but contradictory to other reports in terms of PTZ effects (Auerbach et al. 2013; Beig et al. 2009). Since our experiments were done with Xenopus in anesthesia, the effects of anesthetic drugs should be investigated. Even though we had a control group to investigate the inhibition effects of MS-222 to heart rate variation, the drug-to-drug interactions have not been elucidated. Hence, there is a demand for awake electrophysiology monitoring in drug screening studies. Due to the limited number of Xenopus inspected, there were no general conclusions on how the PTZ influenced the heart and ECG. As many recorded Xenopus ECG signals exhibited abnormalities, PTZ induced negative effects on Xenopus hearts, but the specific influenced locations and mechanism remain uncertain.

### 4.3. Pros and cons of using adult Xenopus in drug screenings

Xenopus is a popular animal model, especially for developmental biology studies. While adult Xenopus were not as widely used in research as their embryonic counterparts in recent decades, previous adult Xenopus studies have sequenced the complete genome and developed many transgenic models, bolstering the adult model for future drug screening studies. As discussed previously, small animal models possessed many advantages for drug screenings, such as lower costs and easier manipulation. However, small mammalian animals often have high heart rate, increase the difficulties and uncertainties for monitoring ECG alterations caused by drugs. Besides, higher heart rates also led to fast drug absorption and dissipation in body. These concerns encouraged researchers to find alternative models. Zebrafish was chosen by many researchers. However, due to their small size, many challenges were still encountered. *X. laevis* had a similar body size to rats, but their heart rate was much slower (11-52 in our monitoring), giving researchers more possibilities to look further into the morphological changes caused by drugs or chemicals. *X. laevis* is a leading model for regeneration research. In (Marshall et al. 2019), stage-dependent cardiac regeneration in Xenopus was discussed. The metamorphosis of Xenopus was significant for developmental studies, as it may help researchers to determine the origin of heart regeneration ability. Additionally, Xenopus are aquatic animals, allowing for the intake of drugs by cutaneous respiration for easier facilitation of drug screening studies. With high absorbability skins of Xenopus, drug application approaches were not limited to injections. The flattened and streamlined body of Xenopus also eased the development of wearable or implantable monitoring devices. Furthermore, Xenopus are relatively inactive, eliminating the influences of high frequency movements, including motion noises and heart rate changes.

Despite their numerous advantages, there are some disadvantages of using adult *X. laevis*. Firstly, there are currently no standards of Xenopus ECG patterns. The three-chambered Xenopus heart differs from the four-chambered human heart. As a result, the species-specific ECG differences need to be investigated to be translated to human models. Second, their 1-2 year long generation time slows the transgenic lines production (Beck and Slack 2001), which constrains reproduction and genetics studies. Third, due to the fully aquatic life habit, more cautions and concerns are needed for wearable and implantable device development.

## 5. Conclusion and Future Work

In this work, a multichannel ECG-EEG recording system on Xenopus was developed. Although, *X. laevis* has not been widely used for studies involving electrophysiological monitoring, we discovered great potential in using this animal model for environmental monitoring and drug screening. The flexible MEA membrane placed dorsally on Xenopus non-invasively recorded the ECG and EEG simultaneously when the frogs were under anesthesia. There were no difficulties in installing the device. Normally, the recordings started within one minute after the Xenopus was sedated after anesthesia. The PTZ and VPA drug tests validated the feasibility of this system in real time monitoring of ECG and EEG for associated neurological and cardiovascular conditions. Both signals displayed detectable and distinct characteristics during different stages of seizures. With excellent skin permeability and easy-to-obtain electrophysiological signals, Xenopus could serve as a desired model for sensing and monitoring of chemical exposure and environmental changes. Future work includes the development of a miniaturized system on a more flexible substrate, such as parylene C, with embedded microelectronics and wireless communication, so that awake electrophysiology signals can be monitored without interferences from Xenopus behavior.

## Acknowledgement

The authors would like to acknowledge the financial support from the NSF CAREER Award #1917105 (H.C.), the NSF #1936519 (H.C), and the NIH SBIR grant #R44OD024874 (M.P.H.L and H.C.). We thank Lauren Schmiess-Heine and Roger Geertsema for managing the Xenopus facility.

